# High-throughput molecular dynamics-based alchemical free energy calculations for predicting the binding free energy change associated with the common mutations in the spike receptor-binding domain of SARS-CoV-2

**DOI:** 10.1101/2022.03.07.483402

**Authors:** Rajendra Bhadane, Outi M. H. Salo-Ahen

## Abstract

The ongoing pandemic caused by SARS-CoV-2 has gone through various phases. From the initial outbreak the virus has mutated several times, with some lineages showing even stronger infectivity and faster spread than the original virus. Among all the variants, beta, gamma, delta and the latest (omicron) are currently classified as variants of concern (VOC) while the remaining are labelled either as variants of interest (VOI) or variants under monitoring (VUM). In this work, we have focused on the mutations observed in important variants, particularly at the receptor-binding domain (RBD) of the spike protein that is responsible for the interactions with the host ACE2 receptor and binding of antibodies. Studying these mutations is particularly important for understanding the viral infectivity, spread of the disease and for tracking the escape routes of this virus from antibodies. Molecular dynamics (MD) based alchemical free energy calculations have been shown to be very accurate in predicting the free energy change due to a mutation that could have a deleterious or a stabilising effect on the protein itself or its binding affinity to another protein. Here, we investigated the significance of six commonly observed spike RBD mutations on the stability of the spike protein binding to ACE2 by free energy calculations using high throughput MD simulations. For comparison, we also used other (rigorous and non-rigorous) binding free energy prediction methods and compared our results with the experimental data if available. The alchemical free energy-based method consistently predicted the free-energy changes with an accuracy close to ±1.0 kcal/mol when compared with the available experimental values. As per our simulation data the most significant mutations responsible for stabilising the spike RBD interactions with human ACE2 are N501Y and L452R.

## Introduction

In the current pandemic caused by SARS-CoV-2(1), the study of viral mutations can give us understanding on the infectivity, pathogenesis, and drug resistance of the virus as well as insight into vaccination efficiency. Further, it may help tracking immune escape routes and consequent emergence of new diseases. During the second wave of COVID-19, SARS-CoV-2 mutations drew much more attention than those of any other virus studied till date. However, it is important to point out that viral mutation is not an unusual phenomenon. For any organism, the mutation rate is the change in genetic information that passes from one generation to the next. The viral mutation rate involves a process that starts from the viral particle’s attachment to the host, its entry, release of genetic material, replication of viral particles, assembly and release or escape from the host cell(2). Viruses show a higher mutation rate than other organisms due to their short generation time and large population size. The mutation rate of viruses also depends on the type of virus. RNA viruses show higher mutation rates than DNA viruses, e.g., HIV virus shows extremely high mutation rate due to its higher replication rate(3). When HIV and SARS-CoV-2 viruses are compared, despite both being RNA viruses, mutations of SARS-CoV-2 are slower than in HIV. There are also many other factors that could influence the mutation rate of a particular virus. It is also important to understand that not every mutation is beneficial to a virus as most mutations are deleterious, leading to the instability of the viral particles(3). The highly deleterious mutation gets rapidly purged while neutral or mildly deleterious stays. These mild mutations impact the virus phenotype and could be beneficial for pathogenicity, infectivity, transmissibility and/or antigenicity of virus particles(4,5).

Since the first outbreak of SARS-CoV-2 in Wuhan, China, the virus has mutated to various lineages. Among these so-called Pango (Phylogenetic Assignment of Named Global Outbreak)(6) lineages some are classified as variants of concern (VOC), variants of interest (VOI) or variants under monitoring (VUM). According to the simplified nomenclature by the World Health Organisation (WHO), Pango lineages such as B.1.1.7, B.1.351, P.1 and B.1.617.2 are named with the letters of the Greek alphabet, alpha, beta, gamma and delta, respectively(7). Currently (February 2022), the B.1.1.529 lineage (omicron) and its descendants are the dominant variants worldwide. Among the known variants, beta, gamma, delta and the latest (omicron) are VOC as of February 2022 (https://www.ecdc.europa.eu/en/covid-19/variants-concern). Of special interest in the present work are the mutations occurring in the receptor-binding domain (RBD) of the SARS-CoV-2 spike protein(8),(9) since that domain plays a major role in the entry of the virus into the host cell by binding to the human angiotensin-converting enzyme 2 (hACE2). For example, a sub lineage of the alpha variant that was first detected in the U.K in December 2020 carries E484K and N501Y mutations in the spike RBD and is 30-50% more infectious than the original virus(10).

The exponential growth in computational power has facilitated the utilisation of *in silico* methods. Computationally expensive free energy calculations that were previously thought to be impossible can now be performed in a few hours. For an ideal free-energy prediction method, the predictive accuracy should be of the same range as that obtained by experimental methods. In a so-called “*alchemical*” transformation, an amino acid is transformed from one state to another via a non-physical pathway. This transformation can be either reversible or irreversible and can be performed by an equilibrium or non-equilibrium method, respectively(11). The major advantage of this approach is the prediction accuracy that matches that reached in experiments(12).

In this work, we performed alchemical free energy calculations, utilising high throughput molecular dynamics (MD) simulations to predict the binding free energy change (with respect to binding partner ACE2) upon six spike RBD mutations commonly observed in current VOC and early VOI lineages. We performed altogether 24 equilibrium MD simulations and 1200 non-equilibrium MD simulations to ensure exhaustive conformational sampling of each of the mutants. We also carried out comparative studies with less rigorous computational methods and discuss the results in the light of available experimental data and/or other computational studies. The results of this comparative study may shed further understanding on the impact of these mutations on infectivity and spread of SARS-CoV-2. Most importantly, we show that the relatively fast free energy calculations may provide a reasonable alternative for expensive experimental mutation studies.

## Results

### Selection of mutations for the study

As per the Pango lineage definition, till date, there are more than 1200 lineages found for SARS-CoV-2(13). The B.1 lineages with the D614G mutation in the spike protein observed in early-2020 gave rise to more than 20% increase in transmissibility compared to the original strain of SARS-CoV-2 Wuhan, China. Other variants also appeared, including many lineages with E484K as the common mutation that was found to reduce the binding affinity of antibodies(14–18). **Figure 1** shows the rough timeline of VOC, VOI and VUM lineages with prominent mutations in the RBD region of the spike protein.

**Figure 1.**
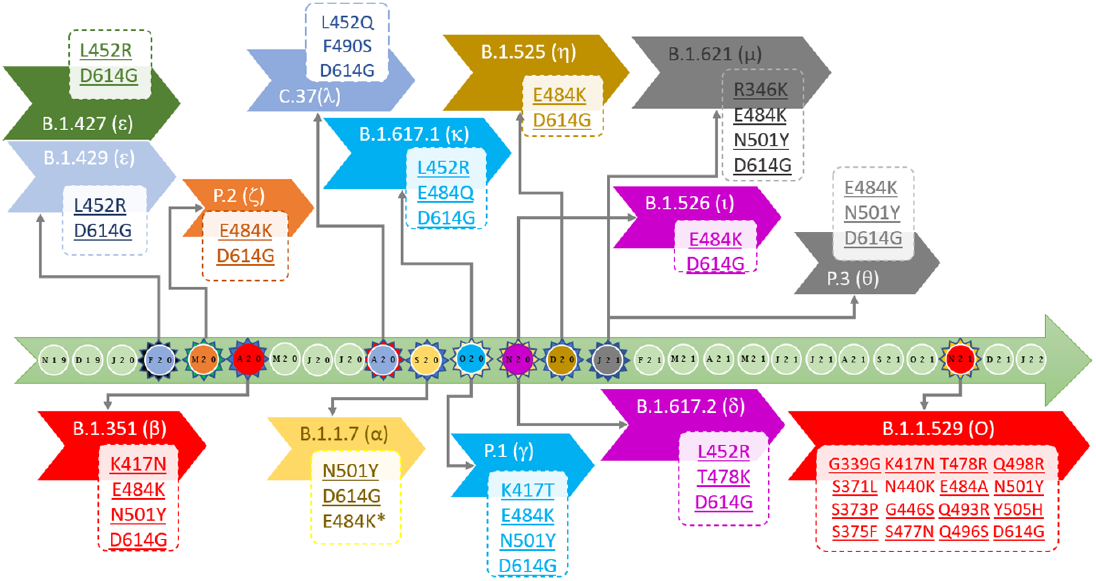
Timeline of various SARS-CoV-2 mutations observed across the Pango lineages. The WHO proposed naming scheme is shown in brackets. The light green central arrow indicates the progress bar, and each circle represents a month, starting from November 2019 to January 2022. The circles with spike-like projections represent the month of the first appearance of new important lineages. Each chevron arrow is for individual lineages; the rectangle boxes contain the information about the key spike RBD mutations (*only in the B.1.1.7 sub lineage). Information on the WHO variants of concern are represented under the central progress bar whereas other variants of interest or under monitoring are above the bar.

The epsilon variant (ε; B.1.427 and B.1.429) with a L452R mutation in the spike RBD was surfaced in California region in the USA in March 2020. The variant showed a 20% increase in transmission,(19) a decrease in susceptibility towards the monoclonal antibody combination containing bamlanivimab and etesevimab as well as reduced neutralization by convalescent and post-vaccination sera(19–21). The P2 variant (zeta, ζ) originated in Brazil during April 2020 and, like epsilon, also showed reduced neutralization by convalescent and post-vaccination sera and monoclonal antibody treatments. At about the same time, South Africa saw a big peak in COVID-19 infections, which was confirmed to be caused by the emergence of the beta variant that carries K417N, E484K and N501Y mutations in the spike RBD. Both K417N and N501Y mutations have been shown to increase the binding affinity of SARS-CoV-2 RBD to hACE2(22). It has been estimated that the beta variant had a 50% increase in transmission rate and a significantly reduced susceptibility towards bamlanivimab and etesevimab monoclonal antibody treatment(23–25). The gamma variant originating from Brazil has also an increased transmissibility and resembles the beta variant but has a K417T mutation instead of K417N. In spring 2021, India saw a big peak in the number of SARS-CoV-2 infections due to the delta variant (B.1.617.2)(26) that has a significantly increased infectivity and viral spread. This variant carries L452R and T478K mutations at RBD. The other variants carry also different combinations of the above-mentioned mutations. The lambda (λ) variant is classified under C.37 lineage and was reported the first time in Peru in December 2020. This variant with L452Q and F490S mutations at RBD has also improved transmissibility and ability to evade vaccine elicited antibodies(27,28), while the following variant (B.1.1.7, alpha) is known for about a 50% higher transmission rate than the original SARS-CoV-2 virus(8). Due to the transmission advantage of the alpha variant and its sub lineage (B1.1.7 + E484K) over B.1 and B.1.1 lineages, it become the dominant variant in the UK in the end of 2020, spreading to the other parts of the world. The kappa variant (B.1.617.1) first detected in India, carries L452R and E484Q mutations and has been shown to have an increased transmutability(29). The mu variant (B.1.621) from Columbia carries an additional R346K mutation not observed in the other variants, and it has shown significant resistance to antibodies elicited by both vaccination and natural SARS-CoV-2 infection(30). The iota and eta variants with E484K mutation at RBD were detected in November-December 2020 in the USA and Nigeria, respectively. In January 2021, the theta variant was reported in the Philippines. It carries E484K and N501Y mutations. Even the latest VOC, omicron, carries some of the same mutations: K417N, T478K and N501Y (+ E484A instead of E484K/Q). The six mutations selected from among the key RBD mutations for this study are R346K, L452R, T478K, E484Q, E484K and N501Y. They include both mutants that have been experimentally shown to affect the spike binding affinity to hACE2 as well as mutants that have either not been studied with respect to the spike-hACE2 binding affinity (to our knowledge) or have been shown to affect for example the immunity (antibody binding) rather than hACE2 binding (**Table 1**).

**Table 1.**
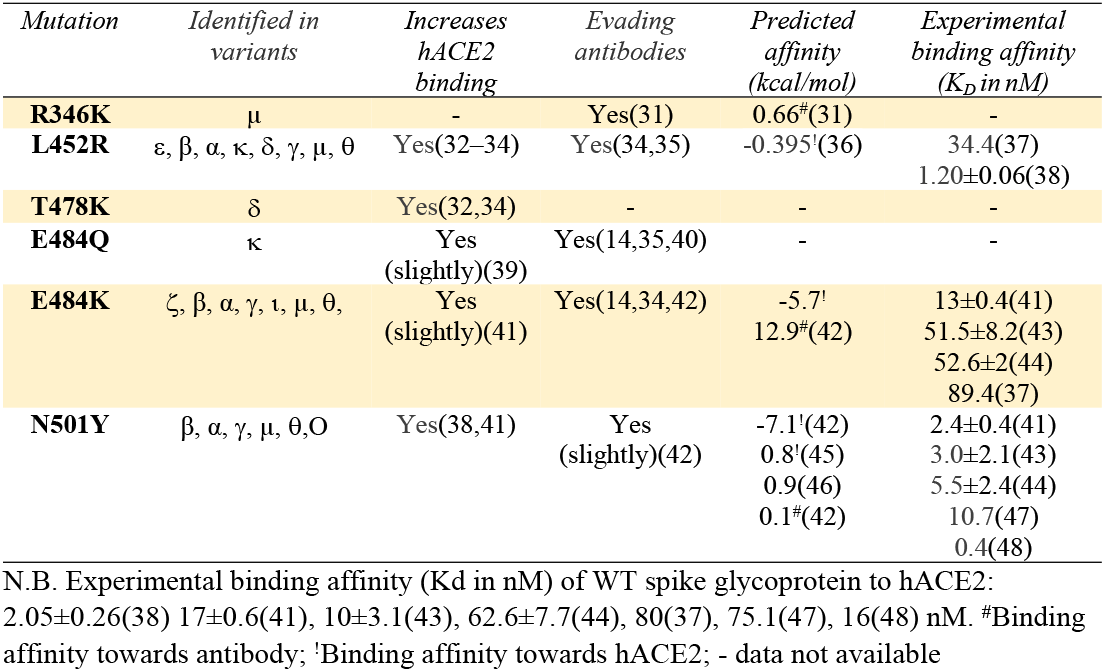
Selected mutations for the study

| *Mutation* | *Identified in variants* | *Increases hACE2 binding* | *Evading antibodies* | *Predicted affinity (kcal/mol)* | *Experimental binding affinity (K<sub>D</sub> in nM)* |
| --- | --- | --- | --- | --- | --- |
| **R346K** | μ | - | Yes(31) | 0.66<sup>#</sup>(31) | - |
| **L452R** | ε, β, α, κ, δ, γ, μ, θ | Yes(32–34) | Yes(34,35) | -0.395<sup>!</sup>(36) | 34.4(37)<br>1.20±0.06(38) |
| **T478K** | δ | Yes(32,34) | - | - | - |
| **E484Q** | κ | Yes<br>(slightly)(39) | Yes(14,35,40) | - | - |
| **E484K** | ζ, β, α, γ, ι, μ, θ, | Yes<br>(slightly)(41) | Yes(14,34,42) | -5.7<sup>!</sup><br>12.9<sup>#</sup>(42) | 13±0.4(41)<br>51.5±8.2(43)<br>52.6±2(44)<br>89.4(37) |
| **N501Y** | β, α, γ, μ, θ, O | Yes(38,41) | Yes<br>(slightly)(42) | -7.1<sup>!</sup>(42)<br>0.8<sup>!</sup>(45)<br>0.9(46)<br>0.1<sup>#</sup>(42) | 2.4±0.4(41)<br>3.0±2.1(43)<br>5.5±2.4(44)<br>10.7(47)<br>0.4(48) |

N.B. Experimental binding affinity (K<sub>D</sub> in nM) of WT spike glycoprotein to hACE2: 2.05±0.26(38) 17±0.6(41), 10±3.1(43), 62.6±7.7(44), 80(37), 75.1(47), 16(48) nM. <sup>#</sup>Binding affinity towards antibody; <sup>!</sup>Binding affinity towards hACE2; - data not available

### Location of the selected mutations at the receptor-binding domain of the spike glycoprotein

The envelope of SARS-CoV-2 has homotrimeric spike glycoproteins that consist of a S1, S2 and a short cytoplasmic domain. To interact with the human angiotensin-converting enzyme 2 (hACE2) the virus utilises the amino acids 331-524 of the S1 domain, that is, the receptor-binding domain (RBD)(49). To structurally analyse the selected mutations, we searched for the available 3D structures of the spike RBD bound with the hACE2 receptor. To date, there are hundreds of SARS-CoV-2 spike protein structures available in the Protein Data Bank (PDB), of which tens of structures are bound with hACE2. This includes structures determined both using X-ray crystallography and cryo-EM. The resolution of these structures varies between 2-5 Å. We set the resolution criteria of 2.0-3.0 Å. As we focused on the RBD of the spike protein, we ignored the structures of full-length spike glycoproteins. We chose the structure deposited by Wang et al. (PDB ID: 6LZG, resolution 2.5 Å)(49) for this study. The characteristic feature of the spike RBD core region (residues 333-437, 507-527) is a twisted, five-stranded β-sheet (formed by antiparallel β1-4 and β7 strands) with short connecting helices (α1-α3) and loops. Between β4 and β7 strands there is an external subdomain (residues 438-506) that is formed by α4 and α5 helices and a flexible loop that connects two short β strands (β5 and β6). This subdomain is also called as the receptor-binding motif (RBM) region since it makes direct contacts with the hACE2 receptor. Except for the R346K mutation that is in the core, right after the α1 helix, all other selected mutations are in the RBM region. L452R is at the β5 strand, T478K and E484K/E484Q mutations are located between the β5 and β6 strands while N501Y precedes the α5 helix(50). Of note, only residues at positions 501 and 484 have direct interactions with hACE2. **Figure 2** shows the location of the mutations selected to be studied here.

**Figure 2.**
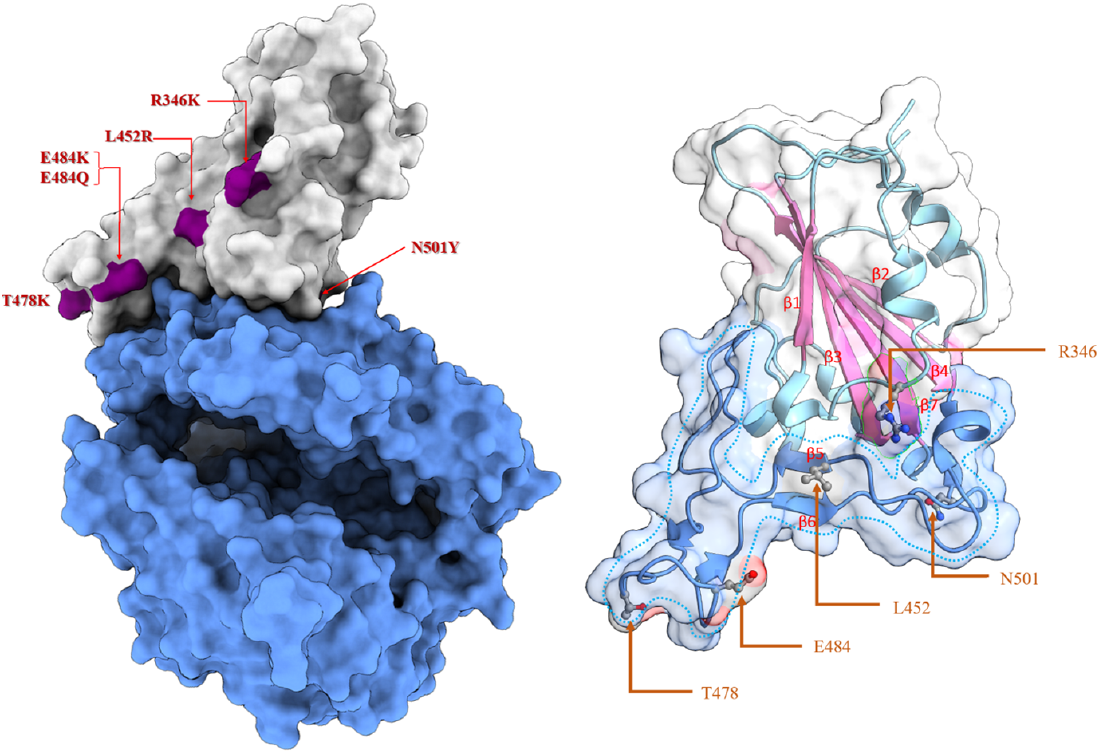
Location of the studied mutations in the spike receptor-binding domain (RBD) of SARS-CoV-2 (PDB ID: 6LZG). Right: Surface representation of the spike RBD in complex with hACE2 (colour scheme: spike RBD in white opaque surface, hACE2 in blue surface, locations of mutations in dark purple colour). Left: Transparent surface and cartoon representation of the spike RBD (colour scheme: the core of spike RBD in transparent white surface, the receptor-binding motif (RBM) domain in blue surface, the core beta sheets are shown in pink cartoon, and the key amino acids in elemental ball and stick; oxygen atoms in red, carbon atoms in grey, nitrogen atoms in blue).

### Prediction of spike RBD-hACE2 interaction stability by non-rigorous methods

Currently, there are various tools available for predicting the functional and/or structural impact of mutations on a protein or protein-protein complexes. In this work, we have utilised Schrödinger’s Prime/MM-GBSA-based residue scanning (Prime, Schrödinger, LLC, New York, NY, 2021)(51,52) as well as CUPSAT(53–55) and FoldX(56–59) tools for predicting stabilising or destabilising effect of the selected mutations on spike RBD-hACE2 complex. In addition, we calculated the effect of the mutations on the stability of the spike RBD alone. The major advantage of these calculations is that they are fast and computationally non-costly. CUPSAT and FoldX tools are also available for free. The downside of these predictions is that they are less accurate and, thus more qualitative than quantitative.

The predicted stabilising and destabilising effects of the selected spike RBD mutations on the RBD binding interaction with hACE2 and the stability of RBD are summarised in Table 2. A ΔΔG value ≥ |1.0| kcal/mol was counted as a significant change (“hotspot”) for CUPSAT/FoldX predictions and ≥ |3.0| kcal/mol for Prime predictions(60). Since all the mutations are found in the viable virus variants, it can be assumed that they should not destabilize the actual spike RDB structure drastically.

**Table 2.**
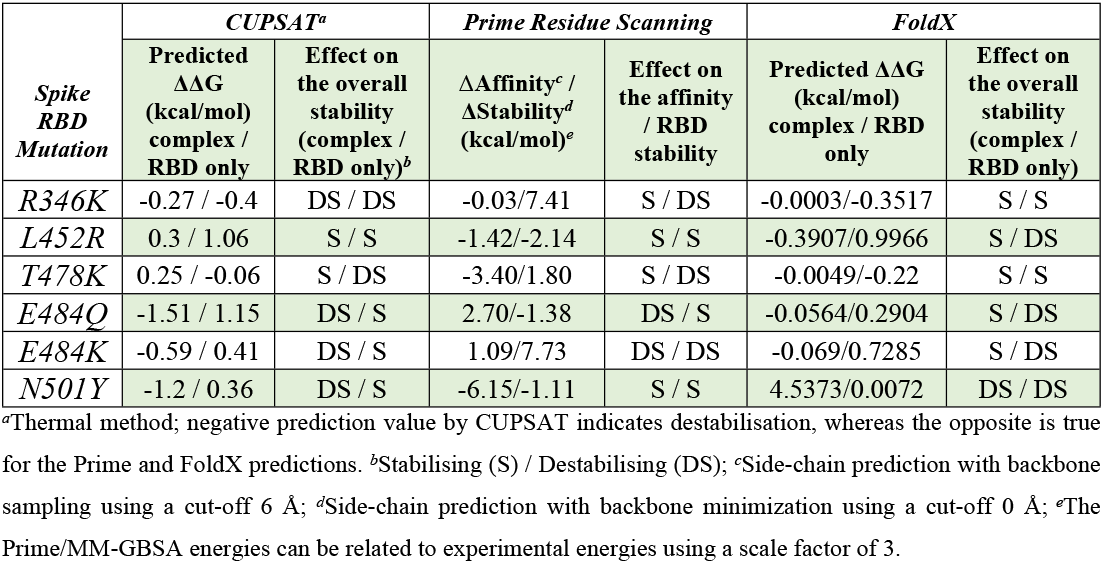
Predicted effect of mutations on the stability of the hACE2-spike RBD complex and the RBD alone

| *Spike RBD Mutation* | *CUPSAT<sup>a</sup>* |  | *Prime Residue Scanning* |  | *FoldX* |  |
| --- | --- | --- | --- | --- | --- | --- |
|  | **Predicted $\Delta\Delta G$ (kcal/mol) complex / RBD only** | **Effect on the overall stability (complex / RBD only)<sup>b</sup>** | **$\Delta\text{Affinity}^c$ / $\Delta\text{Stability}^d$ (kcal/mol)<sup>e</sup>** | **Effect on the affinity / RBD stability** | **Predicted $\Delta\Delta G$ (kcal/mol) complex / RBD only** | **Effect on the overall stability (complex / RBD only)** |
| *R346K* | -0.27 / -0.4 | DS / DS | -0.03/7.41 | S / DS | -0.0003/-0.3517 | S / S |
| *L452R* | 0.3 / 1.06 | S / S | -1.42/-2.14 | S / S | -0.3907/0.9966 | S / DS |
| *T478K* | 0.25 / -0.06 | S / DS | -3.40/1.80 | S / DS | -0.0049/-0.22 | S / S |
| *E484Q* | -1.51 / 1.15 | DS / S | 2.70/-1.38 | DS / S | -0.0564/0.2904 | S / DS |
| *E484K* | -0.59 / 0.41 | DS / S | 1.09/7.73 | DS / DS | -0.069/0.7285 | S / DS |
| *N501Y* | -1.2 / 0.36 | DS / S | -6.15/-1.11 | S / S | 4.5373/0.0072 | DS / DS |

<sup>a</sup>Thermal method; negative prediction value by CUPSAT indicates destabilisation, whereas the opposite is true for the Prime and FoldX predictions. <sup>b</sup>Stabilising (S) / Destabilising (DS); <sup>c</sup>Side-chain prediction with backbone sampling using a cut-off 6 Å; <sup>d</sup>Side-chain prediction with backbone minimization using a cut-off 0 Å; <sup>e</sup>The Prime/MM-GBSA energies can be related to experimental energies using a scale factor of 3.

The prediction tools consistently suggest that mutations L452R and T478K are stabilising the binding interaction. Predictions for the other mutations are inconsistent with these tools. According to the CUPSAT prediction for the spike RBD alone, none of the mutations affects the overall stability of the spike RBD protein significantly (only R346K and T478K are predicted to have a slight destabilising effect while the others are stabilising the protein). The Prime/MM-GBSA-based residue scanning results for some mutants (especially N501Y) were observed to significantly depend on the used refinement cut-off distance for the side-chain predictions. The side-chain prediction with backbone sampling or simple backbone minimization (see Methods) was performed using different cut-off values for including the nearby residues in the refinement. Table 2 shows the affinity change (ΔAffinity) predicted by Prime using the backbone sampling with cut-off of 6 Å. This cut-off gave the most reasonable ΔAffinity value for the N501Y mutant as that mutation is known to increase the spike binding affinity to hACE2 (see the results with other cut-off values in Supporting Information, **Table S1**). On the other hand, the change in spike RBD stability was predicted using minimization of the mutated residue as the refinement method (cut-off 0 Å) to avoid possible errors caused by allowing the protein structure to move more than necessary. According to these results, all but the E484 mutants are predicted to increase the spike RBD binding affinity for hACE2 (T478K and N501Y showing the largest effect on the affinity), while E346K and E484K are predicted to have the greatest effect on the spike RBD stability among the mutants. Of note, the E484K mutant has been experimentally shown to exhibit a slightly lesser thermal stability than the WT whereas N501Y mutant is as stable as the WT(43). Almost all FoldX predictions for the mutants were less than 1.0 kcal/mol in quantity, suggesting an insignificant effect on the hACE2 affinity or spike RBD stability. However, inconsistent with experimental evidence, N501Y was predicted to reduce the spike RBD binding affinity to hACE2 by both FoldX and CUPSAT, and even by Prime with insufficient backbone sampling (**Supporting Information, Table S1**).

### Conventional MD simulation studies

The FoldX optimised structures of the wildtype (WT) and mutant spike RBD bound to hACE2 were used as input for conventional MD simulations. Root mean square deviation (RMSD) and root mean square fluctuation (RMSF) plots are shown in **Figures S1-S2** in Supporting Information. The RMSD of the E484K and R346K mutants converges early, followed by E478K and E484Q and remains stable throughout the MD simulations. More fluctuations in RMSD can be seen for the L452R and N501Y mutants. The fluctuations are prominent towards the C-terminal end of these mutants. The number of intermolecular hydrogen bonds, Coulombic and van der Waals (vdW) interaction energies and the sum of Coulombic and vdW interaction energies for WT and spike RBD mutants with respect to hACE2 were extracted from each 100-ns MD simulation trajectory and averaged (see **Figure 3**).

**Figure 3.**
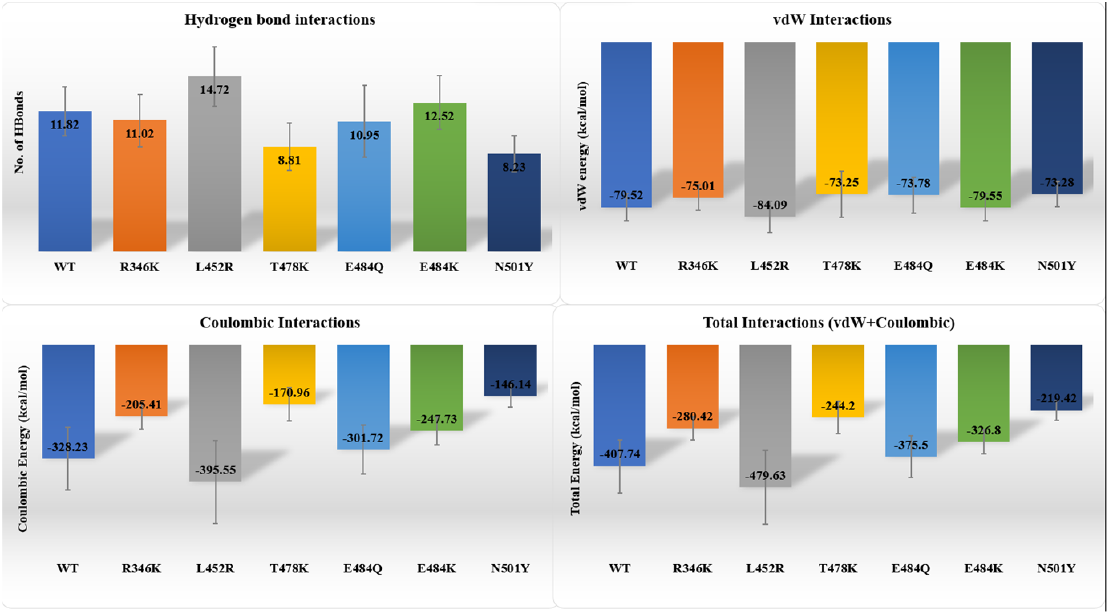
Non-covalent interactions between the SARS-CoV-2 spike receptor-binding domain (wildtype, WT and mutants) and hACE2 averaged over a 100-ns MD simulation.

It was observed that although the N501Y mutation reduces the average number of hydrogen bonding interactions during the simulation, the phenyl ring of the mutant Tyr501 stays very close (within 4.2-6.5 Å) to the phenyl ring of Tyr41 of hACE2 and, thus, is engaged in aromatic stacking interactions throughout the 100-ns MD simulation (**Figures 4a, b**; Supporting Information, **Figures S3 and S4**). In addition, the hydroxyl oxygen of Tyr501 may form a hydrogen bond/a dipole-ion interaction with Lys353 or its phenyl ring engages Lys353 in pi-cation interactions (**Figures 4a, b)**.

**Figure 4.**
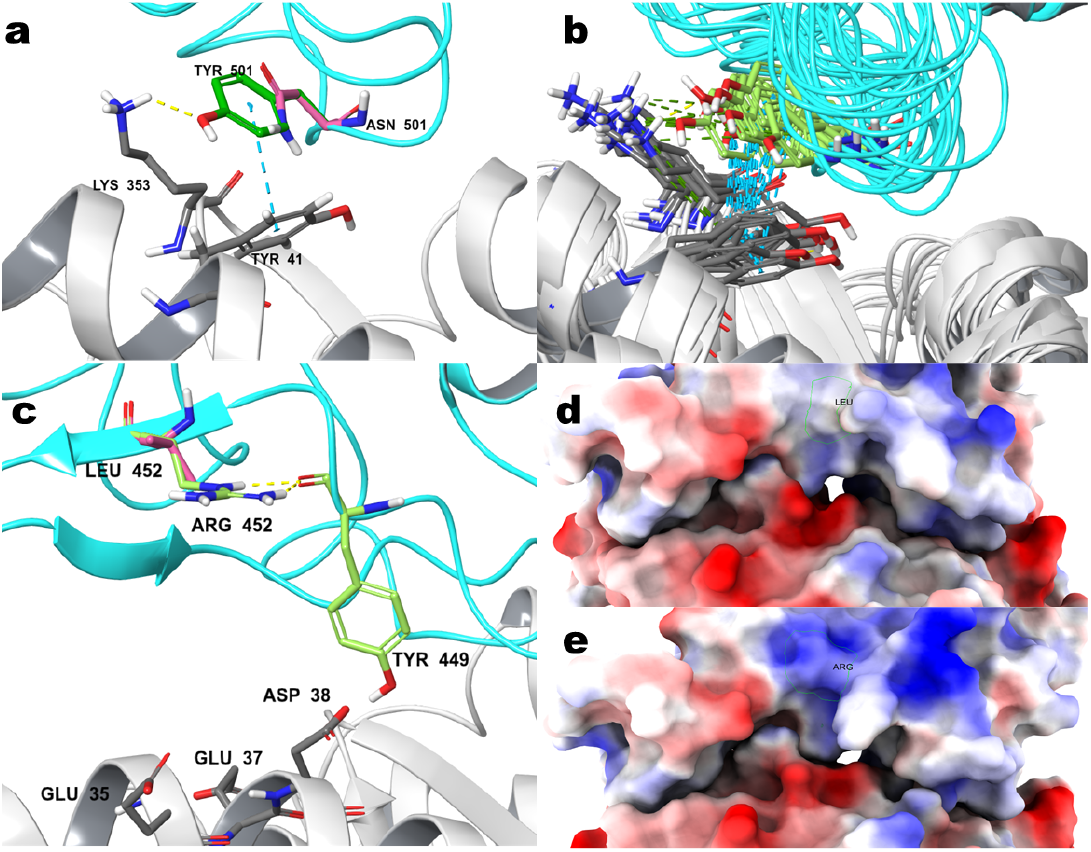
The change in local environment at spike RBD-hACE2 interface caused by N501Y and L452R mutations. **a**. The interactions of Tyr501 with Tyr41 and Lys353 of hACE2 **b**. MD trajectory snapshots taken from the last 50 ns of the 100-ns simulation (each 50^th^ frame) showing the pi-pi stacking between Tyr501 and Tyr41 as well as the pi-cation and hydrogen bond interactions of Tyr501 with Lys353 of hACE2. **c**. A new hydrogen bonding interaction formed between the spike RBD Tyr449 backbone carbonyl and the spike RBD mutant residue Arg452. **d**. Electrostatic surface representation of the spike RBD highlighting the WT residue Leu452. **e**. Electrostatic surface representation of the spike RBD highlighting the mutant residue Arg452. Colour scheme: spike RBD and hACE2 in cyan and white colour cartoon respectively. WT spike RBD, mutated and hACE2 amino acid residues in pink, green and grey colour elemental sticks representation, respectively; hydrogen bond, pi-pi and pi-cation interactions are shown in yellow, cyan and green colour dashed lines, respectively.

Further, the trajectory analysis shows that L452R mutation increases hydrogen bonding as well as Coulombic and vdW interactions. This is likely due to the introduction of the charged residue (Arg452), which causes a change in the local environment (**Figures 4c-4e**), thus favouring increased electrostatic attraction and/or new polar interactions between the proteins. Interestingly, **Figure 4c** shows how Arg452 of the spike RBD mutant interacts with the backbone carbonyl of another RBD residue, Tyr449, thus indirectly providing additional stability to the interaction between Tyr449 and Asp38 of hACE2.

E484K mutation shows a slight increase in H-bonding and vdW interaction energy but the total interaction energy decreased due to the decrease in the Coulombic interaction energy as compared with the WT. The opposite charge seems to be non-optimal for the interaction as the salt bridge between the WT Glu484 and hACE2 Lys31 is lost upon the mutation. In sum, according to the conventional MD simulations, L452R stabilises the spike RBD-hACE2 interactions while all the other studied mutations seem to destabilise the interactions somewhat.

### MD-based alchemical free energy calculations

We did not observe a consensus between the conventional MD simulation results and those from the non-rigorous methods except for L452R. In addition, limited sampling during the MD simulations might affect the ability to predict the stability of the interactions upon mutation. Hence to increase the conformational sampling of the mutants, we carried out high-throughput MD simulations, utilizing non-equilibrium alchemical free energy calculations to investigate the effects of these mutations on hACE2 binding. To perform these calculations, hybrid topology files containing information on both WT and the mutated residues were created.

Among the selected mutations, only R346K and N501Y are non-charge-changing mutations i.e., before and after mutation the change in the charge for these mutants remains equal to zero. This is in contrast to charge-changing mutations (L452R, T478K, E484Q and E484K) where after mutation the net charge change does not equal to zero. Estimating the free energy difference for such charge-changing mutations required an additional step of performing corrections in the charge and was achieved by a so-called analytical post-simulation charge correction method(61). **Table 3** contains the binding free energy calculated before and after the charge corrections (see also **Table S2** in Supporting Information for the uncorrected and corrected binding free energies for each charge-changing mutation in both bound and unbound state).

**Table 3.**
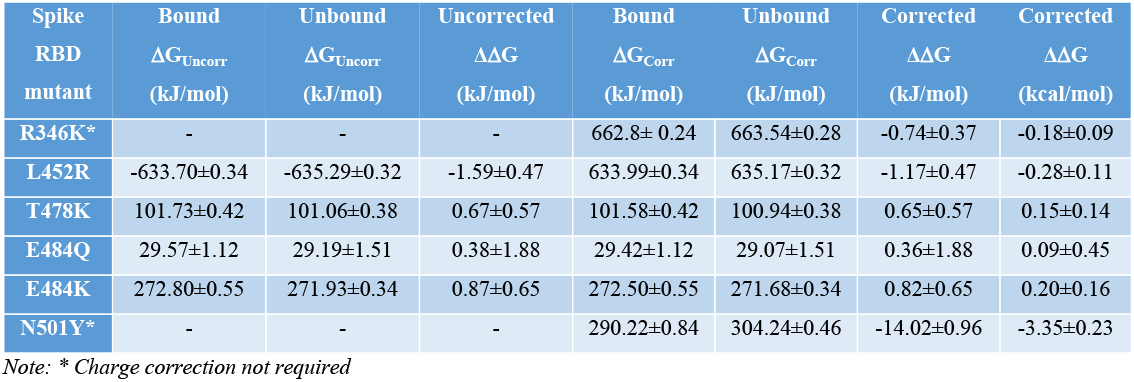
Binding free energies obtained from alchemical free energy calculations for the selected SARS-CoV-2 spike RBD mutations.

| Spike RBD mutant | Bound $\Delta G_{\text{Uncorr}}$ (kJ/mol) | Unbound $\Delta G_{\text{Uncorr}}$ (kJ/mol) | Uncorrected $\Delta\Delta G$ (kJ/mol) | Bound $\Delta G_{\text{Corr}}$ (kJ/mol) | Unbound $\Delta G_{\text{Corr}}$ (kJ/mol) | Corrected $\Delta\Delta G$ (kJ/mol) | Corrected $\Delta\Delta G$ (kcal/mol) |
| --- | --- | --- | --- | --- | --- | --- | --- |
| R346K* | - | - | - | 662.8±0.24 | 663.54±0.28 | -0.74±0.37 | -0.18±0.09 |
| L452R | -633.70±0.34 | -635.29±0.32 | -1.59±0.47 | 633.99±0.34 | 635.17±0.32 | -1.17±0.47 | -0.28±0.11 |
| T478K | 101.73±0.42 | 101.06±0.38 | 0.67±0.57 | 101.58±0.42 | 100.94±0.38 | 0.65±0.57 | 0.15±0.14 |
| E484Q | 29.57±1.12 | 29.19±1.51 | 0.38±1.88 | 29.42±1.12 | 29.07±1.51 | 0.36±1.88 | 0.09±0.45 |
| E484K | 272.80±0.55 | 271.93±0.34 | 0.87±0.65 | 272.50±0.55 | 271.68±0.34 | 0.82±0.65 | 0.20±0.16 |
| N501Y* | - | - | - | 290.22±0.84 | 304.24±0.46 | -14.02±0.96 | -3.35±0.23 |

Note: \* Charge correction not required

According to the free energy calculations, N501Y is the most significant stabilising mutation affecting the binding of spike RBD to hACE2. This is followed by L452R that shows a mild to moderate stabilising effect. The R346K mutation also shows a weak stabilising effect. The remaining three T478K, E484Q and E484K mutations are predicted to be weakly or negligibly destabilising.

We then compared the predicted binding affinity of three mutations with experimentally reported binding affinities (**Figure 5, Table 1, Supporting Information, Table S3**).

**Figure 5.**
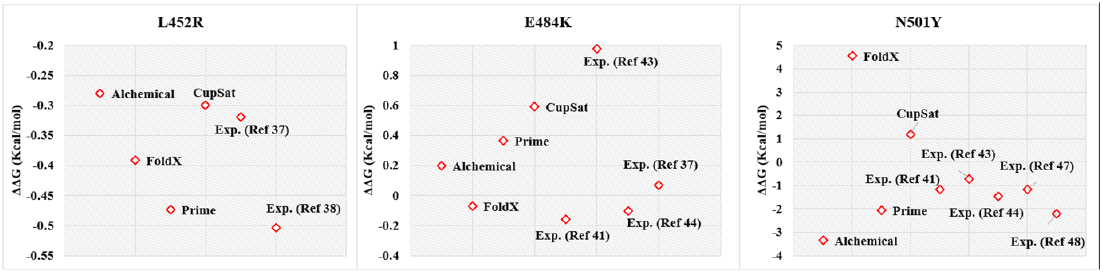
Predicted versus experimentally reported binding affinity of spike receptor-binding domain with hACE2 for L452R, E484K and N501Y mutations. The experimental values were taken from Table 1 and converted to ΔG. The ΔΔG was calculated by subtracting the binding free energy of the WT spike (ΔG_WT_) from that of the mutant spike (ΔG_MUT_). N.B. CUPSAT values (see Table 2) have been converted to correspond to the convention that negative values are stabilizing and positive values destabilising.

**Figure 6.**
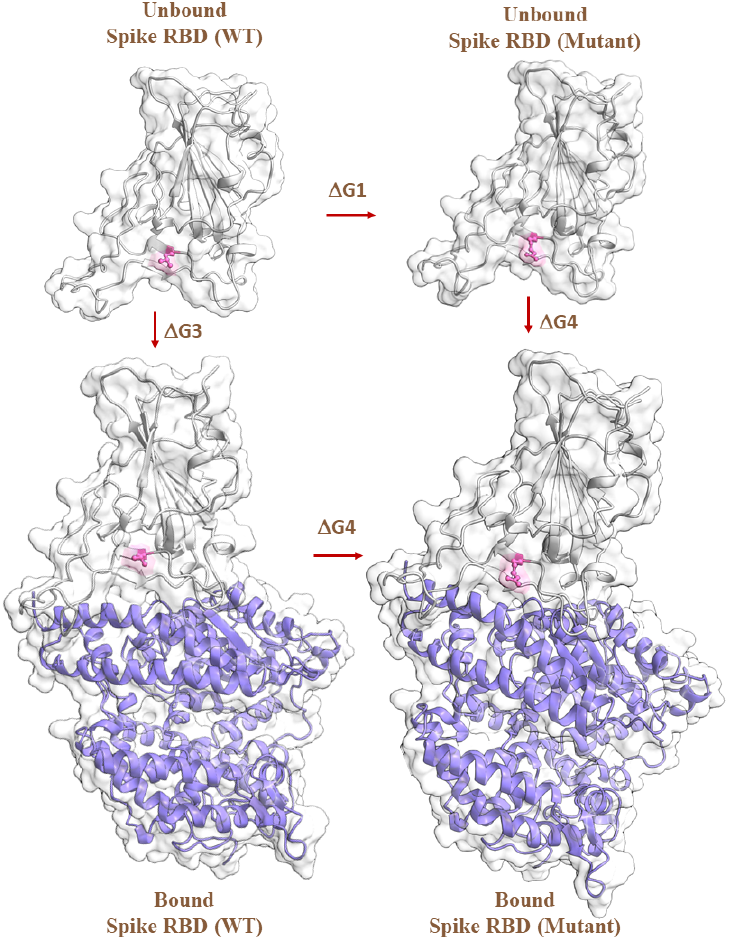
Alchemical free energy cycle represented for the SARS-CoV-2 spike RBD (WT and mutant) in unbound and bound state with hACE2. The arrows indicate the free energy cycle. Leu-452 in WT and Arg-452 in the mutant structure are shown in pink ball and stick representation, spike RBD in grey and hACE2 in purple cartoon with a white transparent surface representation. (The free energy cycle is repeated in same way for all the other mutations.)

The results show that the MD-based alchemical free energy calculations accurately and consistently predicted the affinity when compared with the other prediction methods. The Prime-predicted values also look relatively accurate in Figure 5, but we need to remember that its reliability for some residues may drastically depend on the choice of a suitable distance cut-off during backbone sampling, which here was guided by the available information from experimental studies.

All methods were successful in predicting the effect of the L452R mutation; experimental range of ΔΔG from -0.50 to -0.35 kcal/mol (**Table S1**); prediction range from -0.47 to -0.28 kcal/mol (**Tables 2 and 3**). Even the Prime results with all used refinement cut-offs consistently showed that the mutation is increasing the spike RBD–hACE2 affinity. For the E484K mutant, the corresponding ranges were: experimental ΔΔG from -0.16 to 0.98 kcal/mol and predicted ΔΔG from -0.069 to 0.59 kcal/mol, which suggests a successful prediction even for this mutant that does not have a significant effect on the spike RBD–hACE2 affinity. The Prime-based predictions showed again variability in the values depending on the refinement protocol, some values being negative and some negative (**Table S1**). However, the N501Y mutant was a more difficult task for the non-rigorous methods. The experimental ΔΔG ranges from -0.72 to -2.20 kcal/mol and the predicted ΔΔG from -3.35 to 4.54 kcal/mol, with Prime (cut-off 6 Å) and the alchemical method performing the best and FoldX the worst. (If the other Prime refinement protocols are analysed, by far the worst prediction (7.88 kcal/mol) is given by the backbone minimization with cut-off 0 Å, see **Table S1**).

## Discussion

The viral mutation is an ongoing process and can evolve in response to selective pressures. The SARS-CoV-2 mutation pattern on spike protein observed till date suggests that during the first phase of COVID-19 infections, mutations with increased infectivity overcame the early variant. This can be seen for example from the disappearance of B.1 and B.1.1 lineages and their replacement by more infectious B.1.1.7 lineage in the UK. Though both B.1 and B.1.1.7 carry the characteristic D614G mutation that is known to cause a 20% increase in transmission efficiency, the latter in addition to D614G has the N501Y mutation that ensures more efficient binding of spike RBD to hACE2(62).

Among the six mutations investigated in the current study, only N501Y and E484Q/K are in direct contact with hACE2. Three other mutations are part of the RBM region that forms the binding interface with hACE2. However, none of them has a direct contact with hACE2, while R346K neither has a direct contact nor is part of the RBM region. Consistent with our results, recently reported investigations have shown that L452R and N501Y stabilize the interaction between the spike protein and the hACE2 receptor, thus increasing infectivity(19,36),(63),(64). There are still no reports on the effect of the R346K mutation on the binding interaction with hACE2, but it has been shown that it helps in escaping class 3 antibodies(65,66). The E484K has been shown to have a negligible or small effect on the spike-hACE2 binding affinity (for example, WT Kd 21.3 nM and E484K Kd 19.7 nM(67) (see also Table 1). It has also been identified as an escape mutation that emerges during exposure to monoclonal antibodies (mAbs) C121 and C144(68,69). On other hand there are no other studies on the effect of T478K or E484Q alone on the spike binding affinity to hACE2. As double mutants with L452R they have been shown to increase the binding affinity somewhat(67,70). According to our study, the individual mutations seem not to have a big effect on the spike RBD binding affinity towards hACE2.

When all the three approaches were compared, we observed a consensus only for the L452R mutation. Though we did not observe improved H-bonding, vdW or Coulombic interactions for the N501Y mutant with respect to the WT during the conventional MD simulations, we saw that the phenyl ring of Tyr501 can efficiently form aromatic stacking interactions with Tyr41 and a hydrogen bond or a pi-cation interaction with Lys353 at the hACE2 binding interface. More extensive conformational sampling (i.e., longer MD simulations with several parallel runs) might have given better interaction energies for that mutant, which would also be consistent with the Prime residue scanning results that emphasize the need for sufficient backbone sampling in the environment of the mutation. Notably, the rigorous alchemical free energy calculation method consistently predicted the free energy changes of all mutations with an accuracy close to ±1.0 kcal/mol when compared with the available experimental values.

In sum, the results of this study are consistent with the experimental data on that both N501Y and L452R mutations stabilise the binding interactions between spike RBD and hACE2. On the other hand, neither E346K in the spike RBD core nor the mutations E484Q, E484K and T478K in the RBM offer significant advantage on binding to hACE2. Our study shows also that the MD-based alchemical free-energy calculations work accurately and consistently for different types of mutations (including the more challenging charge-changing mutations) in different structural environments.

## Materials and methods

### Selection of mutants

Based on the reports on increased infectivity of alpha and beta variants and the sudden spike in infections in India in April-May 2021, the key mutations in the spike RBD of these variants were selected for the study.

### Protein Preparation

The 3D structure of the spike receptor binding domain (RBD) that is complexed with the angiotensin-converting enzyme-2 (ACE2) (PDB ID: 6LZG(49)) was retrieved from the Protein Data Bank (PDB)(70). The selected structures were processed using the Protein Preparation Wizard of Maestro (Schrödinger Release 2021-2 New York, NY, USA). Briefly, the missing hydrogen atoms were added to the structure and the hydrogen bond network was optimized using PROPKA at pH 7.0. All water molecules beyond 3 Å were removed and a restrained minimization was conducted using the OPLS4(71) force field with a convergence criterion of 0.3-Å root-mean-square deviation (RMSD) for all the heavy atoms.

### Non-rigorous methods

#### FoldX

The 3D structure of the spike RBD in complex with hACE2 (PDB ID: 6LZG) was stored in the FoldX directory. Using the RepairPDB command, residues having bad torsion angles, steric clashes, or high total energy were identified and repaired. The BuildModel command was then used to create the mutant PDB files and the corresponding WT structures. Finally, using the AnalyzeComplex command the interaction energy between the spike RBD and hACE2 was computed for each mutant and the corresponding WT structure, after which the free energy difference between the WT and the mutants was calculated according to ΔΔG = ΔG_mut_-ΔG_WT_(72).

#### CUPSAT

The CUPSAT (Cologne University Protein Stability Analysis Tool) prediction model uses atom and torsion angle potentials that are specific to the empirical environment of amino acids and were derived from over 4000 protein structures(53). The model’s accuracy to predict the difference in free energy of unfolding between WT and mutant proteins has been shown to be over 80%. The complex stability upon mutation was predicted by providing the PDB ID of the protein-protein complex (6ZLG) to the webserver (http://cupsat.tu-bs.de/) using the thermal experimental method and selecting one mutation at a time. The stability of the spike RBD upon mutation was calculated by providing only the B chain of the PDB structure for the server.

#### Prime Residue Scanning

To predict the effect of individual mutations and to generate input files for MD simulations, residue scanning function of the BioLuminate module in Schrödinger’s Maestro (Schrödinger Release 2021-2: BioLuminate, Schrödinger, LLC, New York, NY, 2021) was utilized(60). It employs the Prime/MM-GBSA (molecular mechanics generalized Born surface area) method (Prime, Schrödinger, LLC, New York, NY, 2021)(51,52) to calculate both the stability and the affinity of the protein in a complex. Using the residue tab, the mutations at selected positions were defined. To see the effect of the cut-off value that defines how far from the mutated residue also the neighboring residues are minimized/sampled, the mutated structure was refined using backbone minimization with cut-off 0 Å and backbone sampling with cut-offs of 0, 4.0, 6.0 and 9.0 Å. The backbone sampling allows adjustments in backbone between the Cα atoms of the adjacent residues, between Cα and Cβ atoms of the residue as well as side chain rotamer adjustments before the final minimization, whereas the minimization adjusts only the side chain after which the whole residue is minimized. The output provides the change in the binding affinity (ΔAffinity) of the specified binding partners (spike RBD binding to hACE2) and change in the stability of the protein (spike RBD) upon mutation (ΔStability) in kcal/mol along with the wildtype (WT) and mutated protein structures. Here, we calculated the stability of the spike RBD separately using just the RBD structure (not the complex) as the input. The output structures from the residue minimization protocol with cut-off 0 Å were used as input for performing the conventional MD simulations. The OPLS4 force field was applied for these calculations.

#### Conventional MD simulations

The WT and mutated structures of spike RBD in complex with hACE2 were submitted to a 100-ns molecular dynamics (MD) simulation using Desmond (Schrödinger Release 2021-2: Desmond Molecular Dynamics System, D. E. Shaw Research, New York, NY, USA, 2021)(73). The simulation systems were prepared using the System Builder tool of the Desmond module. The single point charge (SPC) water(74) was chosen as the explicit solvation model. Each system was neutralized using an appropriate number of Na^+^ or Cl^-^ counter ions. An orthorhombic simulation box, with Periodic Boundary Conditions (PBC) and a 10-Å buffer space between the solute and the box edge, was used for each system. The simulation systems were minimized and equilibrated before the production simulation in a stepwise manner. After the system relaxation, the production simulation was performed in the NPT ensemble for 100 ns, using a reversible reference system propagation algorithm (RESPA) integrator(73). The temperature (300 K) was set using the Nosé–Hoover chain thermostat(75–77), with a relaxation time of 1.0 ps. The pressure was set at 1.01325 bar with the Martyna–Tobias–Klein barostat(78), using isotropic coupling and a relaxation time of 2.0 ps. Long-range interactions were handled using the U-series method(79) and for short range interactions, a cut-off radius of 9.0 Å was used. The MD trajectories were analyzed using the Maestro built-in Simulation Interactions Diagram and simulation event analysis tool and Microsoft Excel360.

#### MD-based alchemical free energy calculations

Alchemical relative free energy calculations exploit and avoid the need to simulate binding and unbinding events by making use of the fact that the free energy is a state function and can be explained as in Equation 1(80,81)).

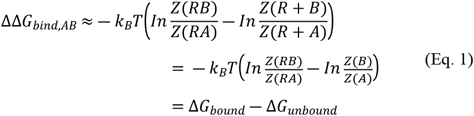

where, Δ*G*_*bound*_ and Δ*G*_*unbound*_ represent the free energy of transformation of state A (WT) to state B (mutant) and the reverse transformation from state B to A. The unbound state is independent on the presence of the receptor in the simulations. The thermodynamic cycle(80) is illustrated in Figure 5.

According to Equation 1 and Figure 3, the difference in the free energy for example for the L452R mutation can be calculated by performing two independent MD simulations and estimating the free energy change from Leu-452 to Arg-452 in the presence or absence of hACE2, i.e., (Δ*G*_*bound*_) and (Δ*G*_*unbound*_), respectively. In an alchemical simulation this change from state A to state B is performed by introducing an alchemical progress parameter, lambda 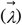 that controls the potential energy function 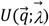 (*q* is the x,y,z coordinates of the simulation system). This is achieved by generating and simulating a hybrid topology of amino acids composed of the atoms that represent both A and B states. A subset of the energetic terms in 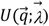 are modulated such that at 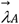 the atoms representing state A are activated and those representing state B are non-interacting “dummy atoms” and vice versa at 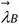. In this way the alchemical transformation is performed and the free energy of mutating A to B in any environment (e.g., bound or unbound state) is computed as per the following Equation 2(80,82).

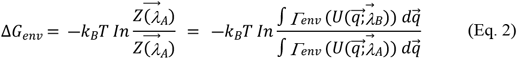

To achieve a good phase-space overlap between the two thermodynamic states, multiple intermediate alchemical states are introduced at values 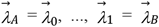 during the simulations. These intermediate states do not belong to either state A or B, but the mixture of the two or a hybrid. The derivatives of the Hamiltonian with respect to λ are recorded at every step and free energies are calculated from the work distributions obtained from integration as per the Equation 3(83).

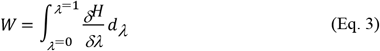

Soft-core potentials are used for both electrostatics and Lennard-Jones interactions as implemented in GROMACS. Finally, ΔΔG is estimated for each mutation by calculating the intersection of the forward and backward work distributions according to the Bennet’s Acceptance Ratio (BAR) method(84).

#### Preparation of Hybrid Topology

The PMX webserver(85,86) was used to generate the hybrid topology files for the selected amino acids. The pdb2gmx command was executed to reassign correct hydrogen atoms in the input coordinate files. The Amber99SB force field was used(12,87) and the output files containing the topology information were generated in a Gromacs readable form.

#### Equilibrium and non-equilibrium MD simulations

All MD simulations (equilibrium and non-equilibrium) were carried out on Puhti HPC cluster(88) with GROMACS-2020.5(89–94). The hybrid topologies generated by the PMX webserver were used as an input. For each mutation, we prepared an individual simulation box and performed forward and backward transition with two simulation systems representing the protein in state A and state B (λ= 0 and λ=1, Figure 3). Amber99SB force field and the Joung and Cheatham ion parameters were used(95). Each state was solvated using a dodecahedral water box with simple point charge 216 (SPC216)(74) 3-point water model and neutralized with Na^+^ and Cl^−^ ions at a 0.15 M concentration. Each simulation system was then energy-minimized for 100000 steps using the steepest descent algorithm(96). This was followed by 1-ns NPT ensemble simulation with positional restraints, using stochastic leap-frog integrator and isotropic Berendsen pressure coupling(97). The final production MD simulations were performed using the stochastic leap-frog integrator for 50 ns(98). All bond lengths were constrained using the LINCS algorithm(99). The constant pressure and temperature condition was maintained using the Parrinello−Rahman pressure coupling(100) at 1 atm and temperature coupling at 298.15 K. A 2-fs time step was used, and the trajectory was recorded at every 10 ps. To treat the long-range electrostatics, Particle Mesh Ewald (PME) method with a direct space cut-off of 1.1 nm and Fourier grid spacing of 0.12 nm were used(101,102). The relative strength of the Ewald-shifted direct potential was set to 10^−5^. Van der Waals interactions were smoothly switched off between 1.0 and 1.1 nm.

To perform the alchemical transformations after the equilibrium MD simulations, fast-growth non-equilibrium simulations were performed to estimate the binding free energy (ΔΔG) value(12,82,83). For this, each equilibrium simulation was used as input for a corresponding non-equilibrium run; the first 25 ns of the trajectory were omitted, and the last 25 ns of the simulation were used to generate 50 snapshots from every 100 ps. The non-equilibrium simulation was performed for a total of 5 ns with the lambda value changing continuously in forward and reverse direction from state 0 to 1 and vice-versa at each time step with a frequency of transition of 2×10^−7^. The standard errors of the ΔG estimates were obtained by bootstrapping and reflect the variability in the datasets (trajectories) analyzed.

#### Correction of artefacts in charge transformative mutations

Among the six mutations studied here, four mutations are charge transformative, which means that when alchemical transformation takes place from state A to state B, the total charge present on the amino acid changes. As we use the hybrid topology approach for the alchemical transformation, during the neutralisation stage only one state (state A or B) is sufficiently neutralised, leaving the system with a net charge of the other state(82). This gives rise to a technical error during the simulation when using the state-of-the-art PME method for handling the Coulombic interactions. Hence, for a non-neutral system, any extra charge is neutralized by the implicit introduction of a uniform background charge(101,102). Due to the contribution of this background charge in the free energy difference, an additional correction scheme is needed to correct the finite charge effect(61,103). Here, we used an analytical post-simulation correction scheme described by Rocklin et al.,(61) which involves an elimination of the following finite size errors; periodicity-induced net charge correction (ΔG_NET_), periodicity-induced undersolvation (ΔG_USV_), residual integrated potential effect (ΔG_RIP_) and correction for discrete solvent effect (ΔG_DSC_). The term ΔG_RIP_ corrects for the residual integrated potential (RIP) and is important if two species are involved polyatomic charge distribution. Hence, for single point charges, a correction scheme utilising the terms ΔG_NET_ and ΔG_USV_ is sufficient to correct the finite charge effect due to neutralisation. Recently, Chen et.al.,(104) compared different approaches required for charge corrections. They combined the (ΔG_NET_) and (ΔG_USV_) terms as per following Equation 4(104):

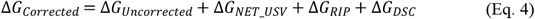

where, ΔG_NET_USV_ term describes both periodicity-induced net charge correction and undersolvation terms. ΔG_RIP_ accounts for the difference between the solute with charges distributed on many atoms and a point charge assumed in Δ*G*_*NET_USV*_ term. ΔG_DSC_ accounts for the difference between explicit water molecules used in the simulation and implicit solvent models considered in the ΔG_NET_USV_ term.

## Author Contributions

**R**.**B:** Performed the MD simulations, the data analysis and wrote manuscript, **O**.**M**.**H**.**S**.**-A:** Supervised the work and critically reviewed the manuscript.

## Acknowledgements

The Sigrid Jusélius Foundation, Biocenter Finland Bioinformatics, CSC IT Center for Science, Tor, Joe and Pentti Borg Memorial Fund and Prof. Mark Johnson and Dr. Jukka Lehtonen are gratefully acknowledged for the excellent computational infrastructure at the Åbo Akademi University.

